# Dot2dot: Accurate Whole-Genome Tandem Repeats Discovery

**DOI:** 10.1101/240937

**Authors:** Loredana M. Genovese, Marco M. Mosca, Marco Pellegrini, Filippo Geraci

## Abstract

The advent of sequencing technologies and the consequent computational analysis of genomes has confirmed the evidence that DNA sequences contain a relevant amount of repetitions. A particularly important category of repeating sequences is that of *tandem repeats* (TRs). TRs are short, almost identical sequences that lie adjacent to each other. The abundance of TRs in eukaryotic genomes has suggested that they play a role in many cellular processes and, indeed, are also involved in the onset and progress of several genetic disorders.

Building upon the idea that similar sequences can be easily displayed using graphical methods, we formalized the structure that TRs induce in dot plot matrices where a sequence is compared with itself. We further observed that a compact representation of these matrices can be built and searched in linear time in the size of the input sequence. Exploiting this observation, we developed an algorithm fast enough to be suitable for whole-genome discovery of tandem repeats.

We compared our algorithm with seven state of the art methods using as a gold standard five collections of tandem repeats: pathology-linked, forensic, for population analysis, genealogic-oriented, and variable TRs in regulatory regions. In addition, we run our algorithm on seven reference genomes to test the suitability of our approach for whole-genome analysis. Experiments show that our method: is always more accurate than the other methods, and completes the analysis of the biggest available reference genome in about one day running at a rate of 0.98Gbp/h on a standard workstation.

## Introduction

A tandem repeat (TR) consists in a certain number of copies of a (typically small) motif sequence that occur adjacent to each other. More realistic definitions of TRs admit a certain degree of heterogeneity among each copy of the motif sequence (in terms of insertions, deletions and mismatches) as well as small insertions between consecutive copies of the motif sequence. The abundance of these repetitive structures in eukaryotic DNAs [1] has been observed since the first sequencing data became available in the early 90s. Although the phenomenon of TRs differential distribution along eukaryotic genomes is not completely understood yet [2], it suggests that TRs are involved in several cellular processes and they modify gene expression. The importance of TRs in these processes has been confirmed in previous studies where the alteration of these sequences has been associated with the onset of several genetic diseases [3, 4].

The steady growth of the number of genetic disorders related to the expansion of TRs has kindled the hope of associating tandem repeats polymorphism with the aetiology of those genetic diseases that are still unexplained. This trend drove the bioinformatics community to focus on research projects aimed at a large-scale analysis of repetitions. Unfortunately, validating a new relevant tandem repeat can be as difficult as finding a needle in the haystack and the success of these projects heavily depends on the sensitivity of the searching algorithms. The difficulty of capturing the variability of satellites and microsatellites into a single comprehensive computational model has encouraged researchers to design new methods for large-scale tandem repeat discovery. Nevertheless, an agreed computational model for non-naive biologically relevant tandem repeats is still far away.

According to different searching strategies and definitions of tandem repeats, several algorithm families have been proposed. Dictionary-based methods [5, 6, 7, 8, 9] leverage on a predetermined set of seeds that is searched along the input sequence and subsequently expanded. The main advantage of this class of algorithms is their efficiency in those situations where the user is interested in a specific, relatively small, class of repeats. The weakness, instead, stands on the fact that an exhaustive search of the tandem repeats is unaffordable because of the exponential relationship between the range of allowed motif lengths and the dictionary size. The dual approach is that of *ab initio* methods [10, 11, 12, 13, 14, 15, 16, 17, 18, 19, 20, 21] that do not require any pre-existing knowledge about the input sequence. Although more complex, this class of algorithms is the most studied and used in practice.

The definition of the divergence (distance) between two units of a motif and the possible presence of an insertion between them is one of the main drivers in the development of tandem repeats discovery algorithms.

The natural presence of insertions and deletions in the genomes has directed many researchers to develop algorithms whose distance is based on the Needleman-Wunsch sequence alignment score described in [22]. The high quadratic cost of this alignment procedure, however, has convinced researchers to shift to algorithms that work in Hamming distance (i.e. the distance is defined as the number of mismatches between two sequences). Hybrid approaches use Hamming distance instead of sequence alignment but allow insertions between consecutive copies of the motif.

Output filtering is a desirable but not mandatory feature of tandem repeats searching methods. Its main advantage is the elimination/reduction of redundancy often caused by software artifacts. Aggressive filtering, however, can cause the removal of relevant results and, as a consequence, the reduction of the algorithm accuracy.

Two distinguishing features have recently gained importance: the ability of managing multi-sequence files (enabling the analysis of reads coming from NGS sequencing) and the possibility of scanning very long sequences (as long as an entire chromosome). Both these features can cause the algorithm to deal with a potentially huge amount of data and thus require fast algorithms that avoid computationally expensive operations, but do not sacrifice output quality.

In this paper we present *Dot2dot*, a novel algorithm for tandem repeat discovery. Our method borrows some ideas from a widely used tool to visually display local alignments between pairs of sequences, namely *dot-plots*. In particular, we observed that aligning a sequence with itself, tandem repeats form a regular pattern. Our algorithm mimics such visual search for these patterns to accomplish tandem repeat discovery. One of the main novelties of our approach is a compact representation of the dot-plot matrices that: 1) allows us to scale at genome-wide analysis, and 2) can find application to other problems where dot-plots are used. Our algorithm belongs to the class of ab initio methods, it allows both a tunable degree of divergence from the consensus sequence and a small insertion between two consecutive motifs. Both fasta and fastq are accepted as input allowing the analysis of NGS sequences. *Dot2dot* implements an optional filtering phase to filter out more aggressively results potentially irrelevant from the biological point of view, and to control the degree of overlap among TRs in the output list. Under the sensible assumption that the longest tandem repeat in the input data is much shorter (by orders of magnitude) than the length of the input sequence, our algorithm runs in linear time (i.e. the running time grows as fast as the input sequence length), thus enabling the analysis at whole genome scale. Besides designing a new searching algorithm, we built five testing datasets covering diverse applicative areas. In particular: we collected from several public sources a set of 45 validated pathology-linked tandem repeats; we compiled the list of coordinates of the CODIS loci including the 7 loci that have been added since January 2017; we mapped on the hg38 reference genome a set of 620 manually-annotated TRs reported in the Marshfield panel of variable loci; and we computed a catalogue of 14, 814 tandem repeats located in upstream regulatory regions. Our test collection and algorithm validation methodology can hopefully become a stimulus to facilitate future research in this field.

## Material and methods

Our algorithm leverages on a data structure at the base of dot plots. Dot plots are used to gain a visual insight of local alignments between two different sequences or even a sequence against itself. Matches between two elements are represented as (typically black) spots. A natural extension of this visual representation of alignments allows the use of color graduation to represent degree of similarity between pairs of elements [23]. The underlying data structure (called dot matrix) is a matrix where the element *M* [*i, j*] stores the degree of similarity between the character in position *i* of the first sequence and the element in position *j* of the other string. When a sequence *s* is aligned with itself, *M*[*i, j*] stores the degree of similarity between the element in position *i* and that in position *j* of *s*.

We observed that tandem repeats form a distinctive pattern on self sequence alignment dot plots and, in turn, this pattern reflects on the underlying dot matrix. Consider for example a pure tandem repeat, since each instance of the motif perfectly aligns with the first instance, it will form a diagonal on the dot plot. All these diagonals will lie stacked over the main diagonal. Counting the number of stacked diagonals we compute the number of copies of the consensus sequence, while from the length of the diagonals we derive the motif length. Fuzziness of tandem repeats can easily be captured within this model. In fact, mismatches correspond to gaps in the diagonals, deletions cause interrupted diagonals and insertions cause the shift of the remaining part of the tandem repeat. Figure 1 shows an example tandem repeat with gaps and an insertion between two copies of the repeated motif. We noticed that the presence in the dot matrix of the above described pattern is a necessary and sufficient condition for the existence of a tandemly repeated sequence in the input, thus an algorithm that locates all and only these patterns ensures a computationally correct and complete tool for tandem repeats discovery.

**Figure 1.**
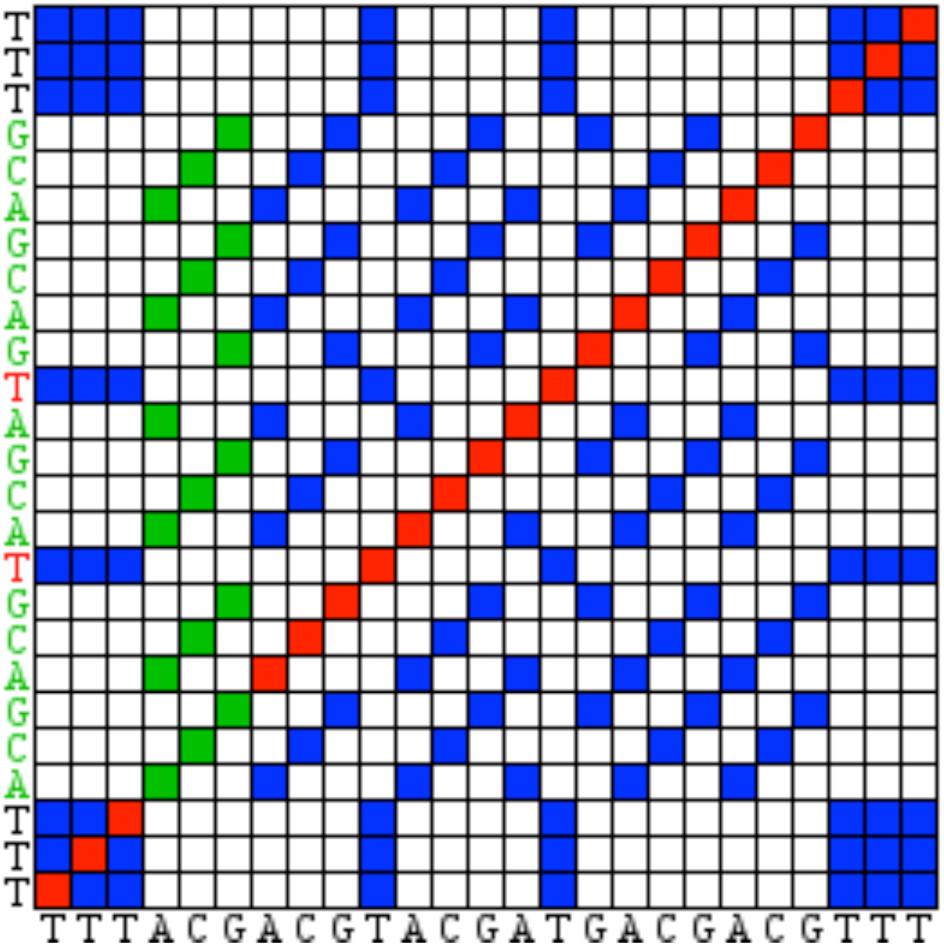
A sample dot matrix of a sequence containing a TR with six copies of the motif ACG with a mismatch in the forth copy and a gap between the second and the third copy. The matrix shows in blue the matching positions, the structure induced by the TR is highlighted in green and the main diagonal is in red. The sequence is reported in both axes and the TR as well as the mismatching positions is highlighted in the vertical axe.

The naive quadratic cost of building, storing and searching of the dot matrix is inadequate for whole-genome analysis. In order to lower the memory consumption and speed up the computation we propose an alternative representation of this data structure that can be built and stored in linear time/space. We also propose a fast searching heuristic algorithm to enumerate all the instances of the tandem repeat pattern in the dot matrix.

### Data structure

In this section we provide some details on how to infer the dot matrix without explicitly building it. For the sake of simplicity we describe here the case of a binary matrix (i.e. where score 1 corresponds to matching characters and score 0 corresponds to mismatches). Indeed, the same procedure extends to the general case without any modifications.

Let *S* = *s*_1_*s*_2_ *…s_n_* be a sequence of length *n* using a finite alphabet Σ (where Σ= {*A, C, G, T*} in our case). Given a character *x* ∈ Σ we define *P* (*x*) as the set of positions of *S* where *s_i_* = *x*.

The comparison of the character *x* and the sequence *S* induces a row *M*[*x*] of *M* containing all zeros except the positions in *P*(*x*) which have score 1. Notice that the content of *M*[*x*] does not depend on the position of *x* in the sequence but only on the content of *S*. As a consequence, comparing the sequence *S* with itself, the resulting matrix *M* has |*P* (*x*)| rows identical to *M*[*x*].

Following the above observation, building the whole *M* it suffices to compute only the vector *M*[*x*] for each *x* ∈ Σ. In order to obtain a direct positional access to the induced dot matrix we build an auxiliary vector *V* of length *n* where in position *i w*e store a reference to *M*[*s_i_*]. Figure 2 shows the data structure corresponding to the matrix depicted in figure 1.

**Figure 2.**
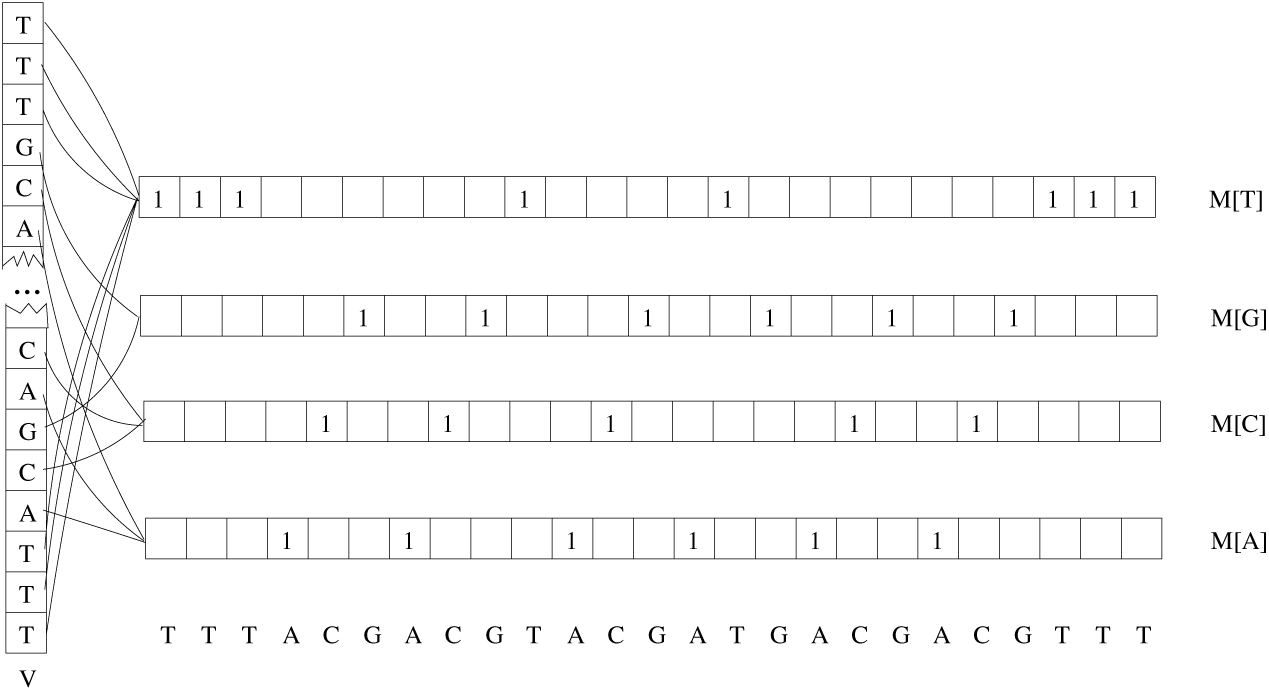
Sample data structure for the matrix in figure 1

Since |Σ| is a constant much smaller than *n*, building the data structure inducing *M* has time/space cost linearly proportional to *n*. In fact, both *V* and the *M*[*x*]*s* are vectors of size *n* that can be filled with a single scan of the sequence *S*.

### TR searching procedure

Searching is a relatively simple procedure that scrolls a window of fixed length along the sequence and attempts to identify for each position whether there are stacked diagonals.

Let *D*(*i, j, l*) be a *l*-long sequence of adjacent cells along *M* starting from position (*i, j*) such that two consecutive elements have coordinates that differ by 1 in terms of both rows and columns and let *W* (*i, j, l*) be the summation of the scores over *D*(*i, j, l*). *D*(*i, j, l*) is a diagonal if and only if *W* (*i, j, l*) *>l* − *δ* where *δ* ∈ [0*, l*] is a user-defined threshold. Given a certain position *i* on the sequence *S*, the event *W* (*i, i* + *l, l*)= *l* corresponds to a pure tandem repeat of motif length *l* and copy number at least 2 starting from position *i*. Again *W* (*i, i* + *l, l*)= *W* (*i, i* +2*l, l*)= *l* corresponds to a pure TR with copy number at least equals to 3 and so on. The parameter *δ* is used to control the degree of fuzziness of tandem repeats. In fact, the event *W* (*i, i* + *l, l*)= *l* − *δ* means that the second copy of the repeat contains *δ* mismatches. Our searching procedure scans the sequence *S* and checks for different values of *l* if the event *W* (*i, i* + *l, l*) ≥ *l* − *δ* is verified. When this happens, our algorithm iteratively attempts to locate further diagonals in positions *i*+2*l, i*+3*l* and so on. The procedure stops when it found an integer *c* such that *W* (*i, i*+*cl, l*) *<l*−*δ*. The sequence *S*[*i, i*+*cl*] is reported as a tandem repeat with motif length *l* and copy number *c*.

We further refined our procedure to deal with insertions and deletions. We restrict to the most significant class of these variants. In particular, insertions can occur only between two copies of the motif sequence and their length is limited to be lower than the motif length *l*. In addition, the length of insertions is fixed within the same TR. This means that if a tandem repeat contains two or more insertions, they must have the same length to be correctly detected. Deletions are modeled as the insertion of a spurious sequence between two copies of the motif string. Although our model of insertions and deletions can appear limited, it has practical advantages and it is consistent with the replication slippage process described in [24]. In fact, without a limit on the length of an insertion nearly every genomic sequence can be confused with a tandem repeat. The constraint on the equality of the insertion lengths within the same tandem repeat helps to predict the correct motif length and copy number when dealing with impure tandem repeats. From the biological point of view, according to the model described in [24], the polymerase is arrested after replicating a unit of the repeat. Then, the realignment between the new strand and the template causes the insertion/deletion to happen between two copies of the TR motif sequence. In presence of an insertion between two copies of the consensus motif the condition *W* (*i, i + kl, l*) >= *l* − *δ* becomes false for a certain *k*. In this case, we seek for a gap checking for a possible diagonal in the next *l* − 1 positions (i.e. *i* + *kl* + 1, *i* + *kl* + 2, etc.). In case of success we take note of the gap and its length and continue the standard searching procedure checking the next diagonal at distance *l*. When the condition on *W* () becomes false again we do not test again all the possible sizes of the gap but we seek only for the length previously annotated.

In terms of asymptotic analysis, the overall computational cost of *Dot2dot* is proportional to the number of times the condition *W* () *>l* − *δ* is tested. A single computation of *W* () takes *O*(*l*) time since it costs *l* accesses to the matrix *M*. Given a certain position *p* ∈ [1, |*S*| − 2*l*] of *S*, there are two cases: either there exists in *S* a TR with *k_p_* copies starting in position *p* or not. In the first case *W* () is computed *k_p_* + *l* times while in the latter case *W* () is computed only once. In general, however, *l* is constant, but *k_p_* cannot be bounded and, thus, it can hold *k* = *max_p_*_∈[1,|_*_S_*_|−2_*_l_*_]_(*k_p_*)= |*S*|*/l* in the worst case. Moreover, a similar condition can hold for every position *p*. As a result, the overall running time of the searching procedure would be |*S*|*lk* = *O*(|*S*|*k*)= *O*(|*S*|_2_) in the worst case.

In terms of average-case analysis, we have to estimate the value 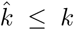 that balances the high cost paid every time a new tandem repeat is found and the low cost paid otherwise. Dealing with real genomic sequences we observed that the probability for a given position to be the starting point of a TR is fairly low. Moreover, the longest TR is order of magnitude shorter than the input sequence. These assumptions would lead to an expected linear running time of our algorithm.

To confirm our hypotheses we estimated the value of 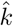 over the entire hg38 reference genome. According to our experiments we measured *k* = 300 and *lk* = 2495. Moreover, the probability of a random position *p* to be the starting coordinate of a tandem repeat is equal to 0.0051. Consequently, the expected value of 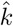 is 1.53.

### Filtering

Exhaustive approaches to tandem repeat discovery suffer from the fact that many reported results may be artifacts rather than proper tandem repeats. For example, finding a tandem repeat of copy number *c*, this class of algorithms return also all the sub-instances of copy number *c*−1, *c*−2, etc. Since this behaviour has also a strongly negative impact on the running time, it would be better avoiding computing these artifacts instead of filtering them a posteriori. In order to solve this problem, we maintain a bit vector to mask those positions that will certainly produce such an undesired result. Once our searching procedure identifies a new tandem repeat it sets the bit corresponding to the first position of each copy of the motif sequence. Marked positions are ignored during the subsequent searching. A second class of artifacts is that of pairs (or sequences) of almost identical tandem repeats whose starting and ending positions differ by only 1 bp. This case arises when the first characters of the TR matches the corresponding characters immediately after the TR. We deal with this class of artifacts allowing TRs having a fractional copy number and removing TRs entirely covered by a longer result. This corresponds to merging together almost identical TRs into a longer result. In particular, when the condition *W* (*i, i + kl, l*) >= *l* − *δ* becomes false (even inserting a gap) we compute the maximum 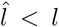 such that 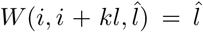 and, 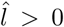, extend the TR accordingly. Notice that we do not allow mismatches in this last copy so that to avoid negative effects on the TR purity.

A last class of artifacts derives from the fact that the same TR can admit several distinct combinations of period length and copy number. This fact is particularly evident for pure tandem repeats. Dealing with impure TRs the only available option is that of computing all the combinations and then evaluating which one better fits a predetermined criterion (in our case we prioritize the purest one). Computing all the possible combinations, instead, is not necessary when dealing with pure TRs. In this case, in fact, testing for multiples of the motif lengh *l* can only produce either TRs of the same size or shorter. In both cases, in absence of a reason to prefer one alternative to another, we return the TR with the smallest *l*, thus we avoid computing the other combinations.

## Results

We experimentally tested our software to assess whether it is able to find non-naive biologically relevant tandem repeats. Moreover, we performed an extensive comparison to evaluate whether using our method can provide an advantage discovering TRs not reported from other methods. As a testing dataset we used five collections of tandem repeats on the hg38 reference genome. In particular we used: 45 disease-related repeats, the 20 extended CODIS repeats, a set of Y-STR loci, 620 markers from the Marshfield panel, and a wide list of tandem repeats located in the regulatory regions. Moreover, we thoroughly scan the literature to find the widest possible pool of alternative algorithms to compare with. Results show that our method is able to find biologically relevant repeats not reported from the other methods.

### Tandem repeats discovery tools selection

Despite the vast literature on tandem repeats discovery tools (see [25] for a recent survey), only few algorithms are usable in practice. In fact, most tools seem to be no longer available or no longer supported [10], [5], [6], [11], [12], [26], [13], [14], [27]; some other is still available but no longer maintained (this is the case for example of [15] that is distributed only in binary form and requires a very old version of the operating system). Some algorithms are subjected to limitations that make comparing with them unfair. In particular: STAR [7] uses a dictionary based approach that makes its computational cost unaffordable even using a small dictionary; MsDetector [16] and IMEx [28] are designed only for microsatellites with motif length less than or equal to 6; the approach in [29] leverages on a specie-specific database; E-TRA [8] can only find perfect tandem repeats; and [30] provides only a visual representation of the distribution of TRs over the input sequence.

At the end of our investigation we identified only seven algorithms that can be realistically employed in daily tandem repeat discovery tasks: TRF [17], mreps [18], tandemSWAN [19], TRStalker [20], SciRoKo [21] TROLL [9] and RepeatMasker [31].

### Parameter setting

All of the considered algorithms require the user to set a certain number of parameters. This procedure in certain cases can bias the outcome of the algorithm and, in turn, its quality assessment. In order to simplify this task most algorithms are provided with a default set of parameters that perform well in most of the cases. Exploring the parameter space of all of the competitor algorithms is beyond the purpose of this paper, thus, after verifying that the default thresholds match the characteristics of our datasets, we always used the default parameters.

### Disease-associated tandem repeats

Due to the proven relationship between repeat expansion and a consistent number of neurological and neuromuscular disorders [3], we tested our algorithm with the aim of evaluating its ability to locate significant tandem repeats associated with diseases. To obtain a convincing list of pathology-related TRs we built our dataset collecting information from several sources. In [4] a list of 44 well-known polymorphic tandem repeats (36 of which confirmed to be associated with a pathology) is given. We removed from this list two entries: SCA31 because the associated repeat is not present in the reference genome since the Spinocerebellar Ataxia type 31 is caused by the insertion of the entire repeat (see [32]), and FSHD (Facioscapulohumeral muscular dystrophy) because the 3.3k D4Z4 macrosatellite repeat is too long for all of the tested algorithms. Due to the lack of the chromosomal coordinates in the above list, we extracted this information from other sources. In a master thesis [33] of the same research group of the article in [4], the authors extend the list including 4 new TRs (two of which linked with a pathology: C9ORF72 and SCA36) and removing two (one linked with the fragile X tremor/ataxia syndrome and one (FRA16A) not directly linked with a specific pathology) providing the coordinates of the TRs in the hg19 reference genome. From the list in [33] we removed 12 elements corresponding to TRs with not confirmed association to a pathology (at the time of this publication) because (as the author mentioned in the thesis) they have been located using the Tandem Repeat Finder software. Since some of the TRs in [4] have now been confirmed as associated with a pathology, and thus their sequence and genomic position is known, we could manually extend our list exploiting *blastn* [34] to locate them.

In particular: according to [35] the sequence of the TR causing the fragile X tremor is a almost pure CGG sequence; in [36] the authors provide the sequence of a repeat expansion in the transcription factor 4 (TCF4) causing the Fuchs corneal dystrophy; in [37] the authors report the motif of a pure TR whose expansion in the noncoding region of C9ORF72 is associated with FTD and ALS; and in [38] the trinucleotide expansion in KCNN3 reported in [39] appears to be associated with schizophrenia. We further extended our list including other notable TRs reported in the literature. In [40] the authors list 29 TRs two of which (one in PHOX2B and one in SOX3) were not included in our list. In [41] an expansion of a pure CGG-repeat in the 5’ UTR of the DIP2B gene is associated with the FRA12A disease. In [42] a mutation of the proneural HASH-1 gene is associated with CCHS. Finally a CAG repeat in POLG1 has been associated with male infertility in [43] and recently with breast cancer risk in [44]. Our final dataset consists in 45 tandem repeats with: a disease-associated polymorphism, period ranging from 3 to 24 bp, and size ranging from 15 to 405 bp.

Although the choice of the version of the reference genome could be considered inconsequential for our purposes, we converted the coordinates of genes and TRs moving to the most recent release of the human genome: hg38. We converted genes manually querying the UCSC genome browser [45] while we used the batch *LiftOver* interface [46] to convert the TRs coordinates. After the conversion we verified that all the TRs sequences were unchanged.

### DNA profiling tandem repeats

Tandem repeats are also largely involved in DNA profiling especially in forensic science. As a result, a large database of short tandem repeats (STRbase [47]) has been made publicly available on the web [48]. Even if the genomic coordinates in a reference genome are not included, this database provides the STR consensus and variants, PCR primers and other useful information to locate each STR. For our testing purposes we used the two most commonly used collections of TR listed in STRbase: CODIS and Y-STR.

#### FBI CODIS

The Combined DNA Index System (CODIS) database consists of 13 tetra-nucleotide tandem repeats spread in 12 chromosomes. As of January 1, 2017 this dataset has been extended with 7 additional tetra-nucleotide TRs: D1S1656, D2S441, D2S1338, D10S1248, D12S391, D19S433, and D22S1045. We collected the coordinates in hg19 of the 13 CODIS core STR loci from the lobSTR [49] website, while we manually retrieved the coordinates of the 7 new tandem repeats. As for the above dataset we converted the coordinates in the hg38 reference genome using the *LiftOver* interface.

#### Y-STR

Y-STR is a collection of short tandem repeats in the *Y* chromosome with period lower than 6 and size ranging from 24 to 166 bp. These repeats are often used for paternity or genealogical tests as well as for forensic purposes. An almost complete list of these loci has been published in [50] and the genomic coordinates in hg19 are available in the lobSTR website. The two markers DYS448 and DYS449 are not TR in a strict sense and consist of two pure STRs interrupted by a small spurious sequence. According to [50], for these marker we included both: the entire STR and the two parts. We completed our list including two extra TRs not mentioned in [50] but present in the lobSTR website: DYS640 and DYS464. The forensic value of DYS464 is studied in [51]. The final collection of Y-STRs tandem repeats consists of 86 loci. As for the above datasets we converted the coordinates in the hg38 reference genome using the *LiftOver* interface.

### Marshfield linkage panel

The Marshfield linkage panel [52] consists of more than six hundred loci distributed across the autosomes containing each a short, highly polymorphic tandem repeat. The main purpose of this panel is that of exploiting the great degree of polymorphism of TRs so as to allow genome-wide population analyses. The original work in [52] neither describes in details the structure of the tandem repeats nor reports their coordinates. Thus, some authors used TRF to pinpoint the repetitive structures. Using TRF is a sensible choice for certain problems (for example in [53] where the authors use the Marshfield panel as a benchmark for genotypization), but it is inappropriate for our purposes.

In the supplementary material of [54], the authors provide the PCR primers, the refseq sequences, as well as a manually-curated description of the tandem repeat structures, for 627 of the Marshfield loci (see The file Pember-ton_AdditionalFile1-11242009 in the the Rosenberg website [55]). Using the refseq sequences as input for *blastn* [34] we located 598 loci. We further obtained the hg38 coordinates of another 22 loci by means of the UCSC’s in silico PCR tool [56]. Finally, we pinpointed the exact coordinates of each TR by a local alignment of the flanking regions of the TR in refseq and the locus in hg38.

The resulting dataset consists in 620 microsatellites with size ranging from 8 to 270 bp.

### Tandem repeats in the regulatory regions

Despite being ubiquitously distributed in eukaryotic genomes, tandem repeats have been reported to be more abundantly present in regulatory regions and in particular in promoters [57], [58]. Their great variability as well as intrinsic instability suggests that mutations of this class of genotypic variation in the promoter regions can influence the observed phenotype [59].

Notwithstanding their importance, a manually curated reference database of variable TRs in promoter regions is not available yet. Partial lists of pure tandem repeats are reported in [60] and [61] while in [62] the authors describe a resource where a partial list is obtained by means of the only TRF software. Besides the database resource, the authors of [62] provide a description of a sensible methodology that we used to recompute the list of TRs in the promoter regions using all the algorithms available to us.

Following the procedure in [62] we downloaded the coordinates of the human genes from the UCSC table browser [63], then we removed non-coding and putative genes as well as genes not in haplotypic regions. After removing duplications we obtained a list of 36096 elements including coding genes and isoforms. In order to ensure that the promoter regions have been entirely covered, as a input sequences we used a 3kb interval from -2kb to +1kb from the transcription start sites. We run all the algorithms on these sequences limiting the TR period to the range from 1bp to 9bp and discarding results with: purity lower than 90% or shorter than 25bp.

Given the reasonably limited number of returned TRs we could refine the results by: re-estimating the correct period length, and trimming spurious endpoints due to the accumulation of errors in the first or last copy of the TR motif. Estimating the period length, we used a brute force procedure that tested all the possible combinations and decided for the shortest value maximizing purity. Then we compared the first and the last instance of each TR motif with the corresponding consensus sequence and trimmed those not matching. Finally we resolved TRs overlapping by removing: the less pure or the TR with higher motif length or the shorter one. At the end of this procedure we obtained a final list of 14, 814 TRs.

### Evaluation metrics

Evaluating the outcome of TR finding algorithms is complicated by a series of choices that can deeply affect the final result. A first issue in comparing the list of repeats returned from an algorithm with the gold standard list is the definition of matching. In the most restrictive setting, one could be interested in the exact match of the starting and ending coordinates, while in general a limited degree of divergence is acceptable. A perfect match is very hard to achieve and it could not be significant because of the subjectivity of the identification procedure. In fact, the coordinates of (more often impure) tandem repeats can slightly differ according to the interpretation given by the curator during the manual annotation phase. In addition, when the flanking regions of a tandem repeat are similar enough to the repeat itself, many alternatives are equally possible. In this case, establishing which is the “correct” repeat, results in an arbitrary choice. This is the case, for example, of the ALS linked repeat in C9ORF72. The repeated sequence consists of three copies of the pure CCCCGG exanucleotide. The annotated tandem repeat is followed by a stretch of four Cs. If an algorithm admits repeats with fractional copy number, it will return the sequence that includes the stretch of Cs. Narrowing the evaluation only to pure repeats does not prevent this problem. In fact, the inclusion of the stretch of Cs does not reduce the purity of the TR. In this case, considering only the exact match and, thus, penalizing the algorithm would be too restrictive.

Another relevant issue is that of dealing with output redundancy (namely, the presence in the algorithm output of overlapping tandem repeats). Although often due to software artifacts, a moderately redundant output may not be problematic when the subsequent analysis is automated, while it could defeat the purpose of an algorithm when only a restricted number of results can be analyzed. An evaluation based on the score of the best hit can penalize those algorithms that employ filtering to reduce the number of biologically irrelevant results.

Finally, the possibility of defining the concepts of true/false positives/negatives, that are at the base of measures like sensitivity and specificity, should be critically examinated. As observed in [64], evaluating an ab-initio algorithm through a gold standard dataset (either the output of an algorithm or a collection of TRs) the true positives are easily defined as the algorithm results that are also listed in the gold standard, while the false negatives are sequences of the testing dataset not reported in the list of results. The problem arises with the other two classes. In particular, it is questionable whether a result reported by the tool but not present in the reference dataset is a false positive or it is a new legal TR that was not previously known. The difficulty of defining the concepts of false positives and true negatives has the effect of hindering the estimation of specificity. Sensitivity, instead, can be computed through the standard formula. However, since sensitivity and specificity are dual evaluation functions that need to be considered as a whole, we decided to use alternative measures. In particular, we used precision and recall.

In order to address the issue of matching predictions and gold standards, we used the Jaccard coefficient as a measure of matching. This coefficient has originally been defined to measure the degree of similarity between sets. Subsequently, it has been extended to measure the overlap between intervals. In short, it is defined as the ratio of the length of the intersection of two intervals divided by the size of their union. The Jaccard coefficient is bounded in the range [0, 1] and to a higher value corresponds a higher degree of matching. Entirely covering a TR is necessary but not sufficient to get the highest score, in fact, the Jaccard coefficient has value 1 only when the comparing intervals have exactly the same coordinates. Investigating several alternative values of Jaccard score as threshold to decide that a repeat has been identified, we obtained a complete picture of the algorithms’ behaviour.

Dealing with issue of redundancy, we used three measures: the average number of results covering an element of the gold standard, the average precision and the average recall. Let *T* =(*t*_1_*,…,t_n_*) be a dataset of tandem repeats, *R* the set of results of a given algorithm, and *R*(*t_i_*) the subset of *R* overlapping with *t_i_* by at least 1bp We further denote *jac*() as the Jaccard coefficient between two genomic intervals. The average precision and average recall are defined as follows:

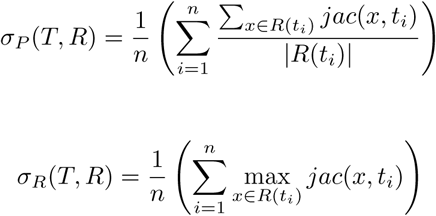

The above measures have several advantages. Firstly, they are independent from the arbitrary choice of a threshold value. Secondly, results over different datasets can be merged into a single score or can be directly compared. Lastly, it is possible to compare algorithms that apply filtering with methods that do not use it. In fact, even in a case where the filtering phase removes the most overlapping repeat causing the Jaccard score of the second best hit to drop under threshold, the removal of a promising repeat could be balanced by the removal of a certain number of irrelevant repeats with low Jaccard score.

## Discussion

In this section we report and discuss details about the outcome of our comparative evaluation. We run our algorithm (*Dot2dot*) using two sets of parameters: one with the default values but without any filtering, and one with the same values but enabling the most stringent filtering. We also set a time limit of two hours to complete a single run over an input sequence.

Figure 3 shows the number of TR identified in each test collection by the compared algorithms for increasing values of the Jaccard score. At a first sight, it is possible to observe that, for two of the profiling datasets: CODIS (fig. 3 (b)) and Y-STR (fig. 3 (c)) our algorithm is always able to find at least as many repeats as the other algorithms independently from both the use of filtering and the threshold of Jaccard score. These datasets highlight the weakness of two of the compared algorithms: TRStalker and TandemSWAN. In fact, although the length of the CODIS and Y-STR input sequences are limited to at most few million base pairs, TRStalker has never been able to complete within the time limit of two hours. TandemSWAN, instead, terminates each run in less than two minutes, but identifies an unexpectedly low number of tandem repeats. This result depends indeed on the (unavoidable) filtering step of the algorithm. We conjecture that the high purity of the CODIS and Y-STR repeats is likely to induce the algorithm to filter out most of the TRs because considered uninteresting.

**Figure 3.**
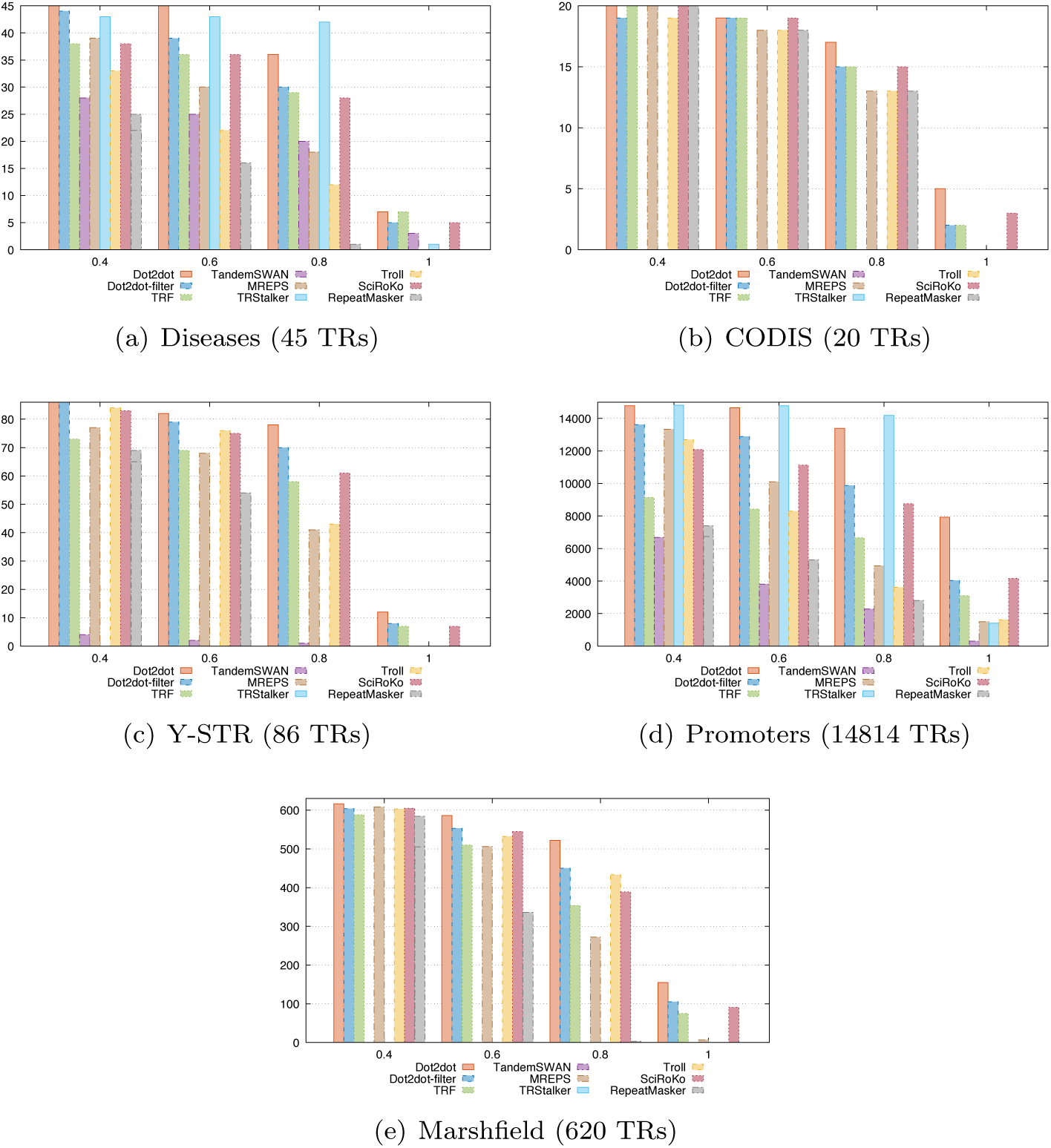
Number of TRs identified by each algorithm for increasing values of the Jaccard coefficient.

We obtained a similar result on the Marshfield panel (fig. 3 (e)). In this case, we used the entire hg38 reference as input so as to perform an evaluation at genomic level. Again, TRStalker did not complete within the limit of two hours, while TandemSWAN located almost none of the tested TRs. Interestingly, comparing fig. 3 (e) with fig. 3 (b) and fig. 3 (c), we saw that the relative ranking of all the algorithms remains unaltered. This fact indicates that the length of the input sequence does not introduce any bias in the behaviour of the tested tools.

According to figure 3 (a) and 3 (d), when dealing with pathology-linked TRs and repeats in the regulatory regions, our algorithm is still at least as accurate as the other algorithms, except for TRStalker that achieve an higher score when the threshold for the Jaccard score is set either to 0.6 or 0.8. This higher accuracy, however, is the effect of a very large redundancy due to the exhaustive enumeration of all the possible alternative TRs covering the same locus. In fact, as table 1 shows (see column RPL), TRStalker covers a locus with an average of over 100 different results.

**Table 1.** Average precision *σ*(*P*), average recall *σ*(*R*), and average number of reported Results covering a target locus (RPL) of the compared algorithms and datasets.

| Algo | Diseases |  |  | CODIS |  |  | Y-STR |  |  | Marshfield |  |  | Promoters |  |  |
| --- | --- | --- | --- | --- | --- | --- | --- | --- | --- | --- | --- | --- | --- | --- | --- |
| | $\sigma(P)$ | $\sigma(R)$ | RPL | $\sigma(P)$ | $\sigma(R)$ | RPL | $\sigma(P)$ | $\sigma(R)$ | RPL | $\sigma(P)$ | $\sigma(R)$ | RPL | $\sigma(P)$ | $\sigma(R)$ | RPL |
| Dot2dot-Filter | **0.830** | 0.835 | **1.02** | **0.860** | 0.860 | **1.00** | **0.877** | 0.879 | 1.03 | **0.856** | 0.860 | **1.03** | **0.825** | 0.803 | **1.01** |
| Dot2dot | 0.678 | 0.899 | 6.04 | 0.721 | **0.899** | 6.35 | 0.706 | **0.916** | 6.11 | 0.637 | **0.905** | 7.09 | 0.667 | **0.934** | 4.80 |
| TRF | 0.722 | 0.763 | 1.43 | 0.836 | 0.850 | 1.10 | 0.770 | 0.786 | 1.06 | 0.784 | 0.794 | 1.10 | 0.663 | 0.548 | 1.37 |
| MREPS | 0.630 | 0.686 | 1.66 | 0.659 | 0.818 | 2.10 | 0.617 | 0.712 | 2.11 | 0.662 | 0.755 | 1.99 | 0.575 | 0.687 | 1.88 |
| TRStalker | 0.464 | **0.901** | 155.88 | — | — | — | — | — | — | — | — | — | 0.422 | 0.929 | 116.06 |
| SWAN | 0.372 | 0.575 | 1.97 | 0 | 0 | 0.00 | 0.021 | 0.029 | 2.50 | 0 | 0 | 0.00 | 0.663 | 0.447 | 2.24 |
| Troll | 0.508 | 0.571 | 4.14 | 0.682 | 0.797 | 3.20 | 0.682 | 0.777 | 2.77 | 0.683 | 0.792 | 3.22 | 0.517 | 0.627 | 2.09 |
| SciRoKo | 0.724 | 0.752 | 1.15 | 0.819 | 0.867 | 1.25 | 0.827 | 0.841 | 1.10 | 0.810 | 0.830 | 1.13 | 0.808 | 0.709 | 1.05 |
| Repeatmasker | 0.452 | 0.475 | 1.17 | 0.812 | 0.822 | 1.05 | 0.721 | 0.729 | **1.02** | 0.767 | 0.775 | 1.05 | 0.743 | 0.432 | 0.58 |

Although the ability of identifying the repeats of the gold standard with a reasonably high Jaccard score is a desirable property, achieving a high recall at the cost of a large output redundancy can greatly reduce the usefulness of an algorithm in practice. In particular, a good filtering strategy should increase the average precision at the cost of a negligible reduction of accuracy. Aimed at quantifying redundancy and evaluating its effects, we measured the average number of results per locus (RPL), the average precision and the average recall of each algorithm for the five test collections.

Table 1 shows that an aggressive filtering is likely to have a positive impact on the average precision as well as on controlling the redundancy of the output. In fact, the algorithms implementing the most aggressive filtering strategy (*Dot2dot-Filter*, TRF and SciRoKo) achieve the highest values of average precision. In contrast, a high level of redundancy is not necessarily a guarantee of a high average recall. This is the case, for example, of Troll that has the third highest number of results per locus but achieves one of the lowest average recall. *Dot2dot* and TRStalker, which have the most verbose output, achieve the highest average recall. However, in spite of having almost the same average recall, *Dot2dot* returns a number of results per locus of the same order of magnitude of the other methods, while TRStalker produces two orders of magnitude more results per locus.

Comparing *Dot2dot* run with and without filtering, we observed how our filtering strategy has a remarkably positive impact in terms of a drastic reduction of overlapping results, passing from about 5 results per locus to approximately 1. Filtering causes an increase of average precision of about 0.15 allowing *Dot2dot* to attain the highest score over all the tested datasets at the cost of a limited reduction of average recall (−0.06). However, this lower recall does not change the relative ranking of our algorithm in the comparison with the other methods and, as shown in figure 3, has little impact in terms of best-hit accuracy.

### Running time

Although running time is a secondary feature for tandem repeats discovery algorithms, it can still have an important role for whole-genome analyses. In fact, as observed before, TRStalker is so computationally demanding that it was unable to complete the analysis of most of the input sequences within two hours, and, thus, it could not run on entire genomes. In order to test the possibility of using the selected algorithms for whole-genome analysis we run them on 7 reference genomes, with sizes ranging between 1.3Mbp and 27Mbp, available on the NCBI website. We used a MacOS-based workstation endowed with a processor Intel Xeon 3.1Ghz and run the software in single thread mode. For the software running only under Linux (namely: tandemSWAN and Troll) we had to use a different hardware. We computed the performance difference between the two machines by comparing the speed of *Dot2dot* over chromosome 1 of hg38. We found that the Linux server is 1.4556709189 times faster. As a result, we used this constant as a correction factor for the running time of tandemSWAN and Troll.

We report in table 2 the running time of the tested algorithms. We excluded TRStalker because it cannot run on these large sequences and Repeat Masker be cause its running time is dominated by the identification of other classes of repetitions different from tandem repeats. Since the computational cost can vary according to the richness of the output, we also evaluated the algorithms in terms of throughput and discovery rate. As a throughput measure we computed the number of results returned per second (figure 4), while the discovery rate is the average number of tandem repeats found per Kbp (figure 5). Since we encourage the use of filtering for whole-genome analysis we run *Dot2dot* only with filtering. However, we notice that this choice does not give *Dot2dot* any advantage in the comparison. In fact, without filtering our software would run slightly faster and would have a much higher throughput and discovery rate (data not shown).

**Table 2.**
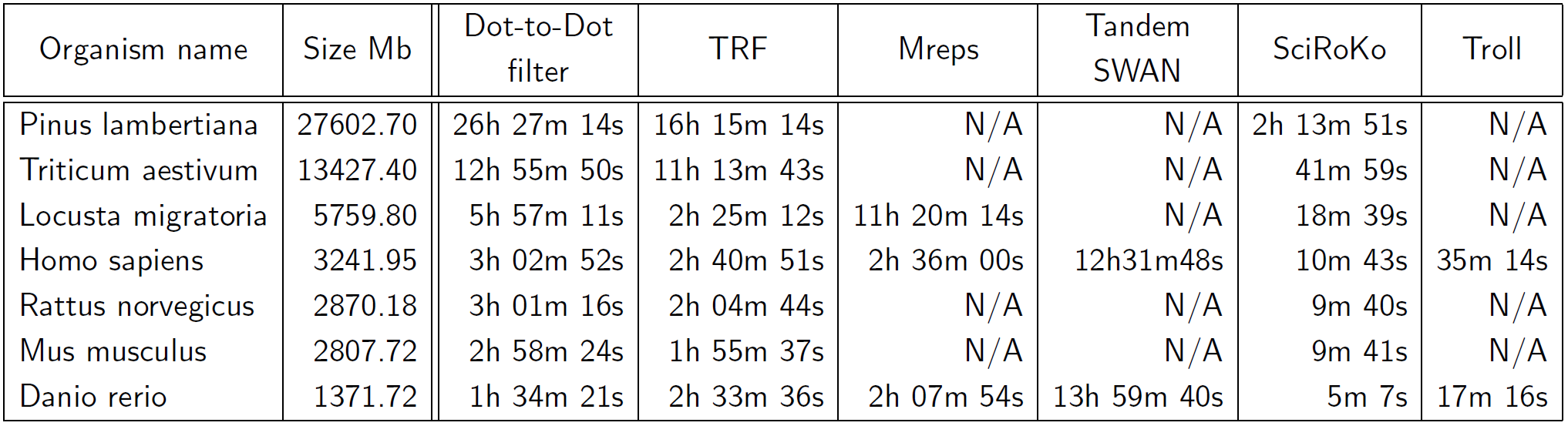
Running time of the compared algorithms over 7 common reference genomes. The genome dimension is measured as the size in Mb of the corresponding fasta file.

| Organism name | Size Mb | Dot-to-Dot filter | TRF | Mreps | Tandem SWAN | SciRoKo | Troll |
| --- | --- | --- | --- | --- | --- | --- | --- |
| Pinus lambertiana | 27602.70 | 26h 27m 14s | 16h 15m 14s | N/A | N/A | 2h 13m 51s | N/A |
| Triticum aestivum | 13427.40 | 12h 55m 50s | 11h 13m 43s | N/A | N/A | 41m 59s | N/A |
| Locusta migratoria | 5759.80 | 5h 57m 11s | 2h 25m 12s | 11h 20m 14s | N/A | 18m 39s | N/A |
| Homo sapiens | 3241.95 | 3h 02m 52s | 2h 40m 51s | 2h 36m 00s | 12h31m48s | 10m 43s | 35m 14s |
| Rattus norvegicus | 2870.18 | 3h 01m 16s | 2h 04m 44s | N/A | N/A | 9m 40s | N/A |
| Mus musculus | 2807.72 | 2h 58m 24s | 1h 55m 37s | N/A | N/A | 9m 41s | N/A |
| Danio rerio | 1371.72 | 1h 34m 21s | 2h 33m 36s | 2h 07m 54s | 13h 59m 40s | 5m 7s | 17m 16s |

**Figure 4.**
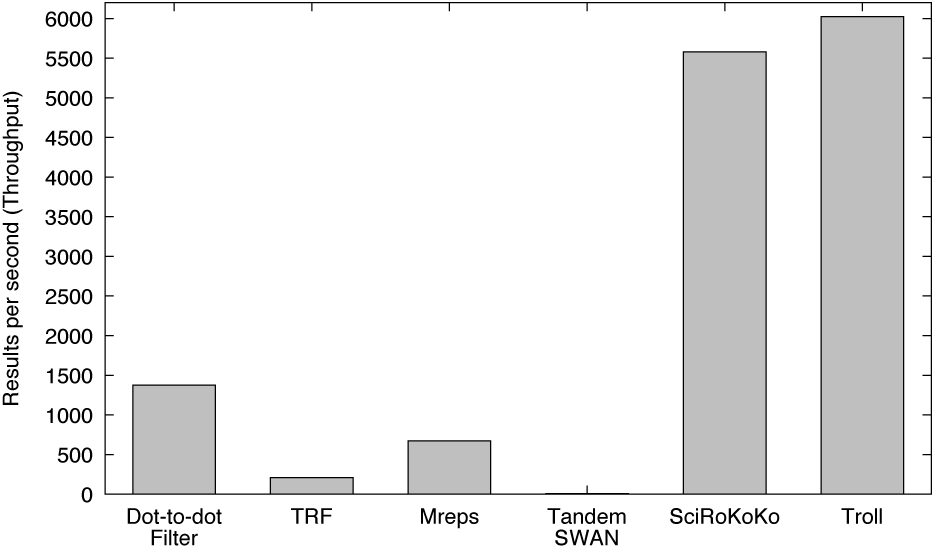
Average number of returned results per seconds (throughput) of the compared algorithms.

**Figure 5.**
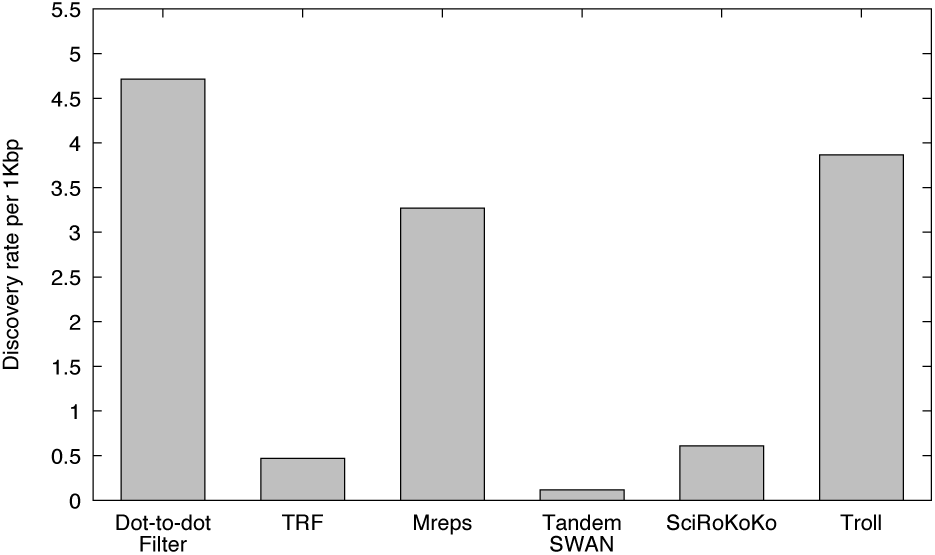
Discovery rate expressed as the expected frequency of reported tandem repeats per sequences 1kbp long.

As table 2 shows, three algorithms fail on some of the genomes. This is mostly due to intrinsic limits hardwired in the software. For example, Mreps has a limitation on the length of the longest consecutive stretch of Ns in a sequence. Because of it, Mreps could not run on relatively small genomes like that of Mus musculus but could run on the bigger genome of the Locusta migratoria instead.

SciRoKo is the fastest algorithm with the second highest throughput, however, the discovery rate is quite low. This result is probably due to the fact that SciRoKo returns TRs with motif length only up to 6bp. Troll is the second fastest software (the first in terms of throughput) with an outstanding discovery rate. However, it has a practical limit on the overall length of the input sequences that should not exceed 2^31^bp (about 2.1Gbp). As a result, in order to run it on the human genome we had to search each chromosome independently and merge results together.

Except for a single case (for the Danio rerio genome), TRF runs faster than *Dot2dot*. The difference is often not high, but it can become consistent for the biggest genomes. However, comparing the two algorithms in terms of throughput, our software finds on average more than 5 times more results per second than TRF (see figure 4). Since *Dot2dot* filtering policy does not allow overlapping results, this gain is not due to the redundancy of the output.

Data on table 2 confirms also the average-case cost analysis of our algorithm. In fact, the *Dot2dot* running time grows proportionally with the genome’s length with a small stable constant factor.

## Conclusion

In this paper we presented *Dot2dot*: a new algorithm for tandem repeat identification in a target genome. Our model of repeat has shown to be general enough to capture well various classes of tandem repeats with different characteristics: pathology-linked, forensic, for population analysis, genealogic-oriented, and repeats in the regulatory regions. Our experiments have shown that *Dot2dot* is fast and effective since it was able to identify almost all the tandem repeats of our test collections with an accuracy of at least 0.7 in terms of Jaccard score. Even applying a severely stringent filtering where overlap among the returned repeats is not allowed, our algorithm has still shown to be more accurate than the alternative tested tools. Our tests over the entire human genome have confirmed the hypothesis that the longest tandem repeat found (with a length of 2495bp) is several order of magnitude shorter than the input sequences and thus our repeat identification and filtering algorithms run in linear time with the input length allowing the analysis of a whole human genome within few hours in a standard workstation.

Software implementing *Dot2dot* is freely available and can be used without any limitations. To make it useful in the daily laboratory practice we enabled it to read standard formats for both assembled genomes (fasta) and NGS data (fastq) as well as return its output also in a standard format (bed).

Another contribution of this paper is that of proposing a rigorous assessment methodology of tandem repeats discovery algorithms as well as providing five reference collections of TRs (four of which manually curated). We hope that the proposed methodology and datasets can help to facilitate future research in this field.

## Competing interests

The authors declare that they have no competing interests.

